# Self-Organization Through Local Cell-Cell Communication Drives Intestinal Epithelial Zonation

**DOI:** 10.1101/2025.11.14.688372

**Authors:** Yael Heyman, Michal Erez, Phil Burnham, Mor Nitzan, Arjun Raj

**Author notes:** Equal contributions (Yael Heyman, Michal Erez).

## Abstract

The intestinal epithelium exhibits zonated gene expression along the crypt-villus axis, with distinct transcriptional programs in enterocytes at the villus top versus bottom. However, the mechanisms establishing these spatial patterns remain unclear. Three models could explain zonation: external gradients, cell-intrinsic temporal programs, or local self-organization. Using spatial transcriptomics and perturbations of two-dimensional intestinal organoids, we show that zonation emerges via spontaneous self-organization without mesenchymal, neural, or vascular inputs. Cell-intrinsic models were eliminated by transplanting cells into established monolayers; transplanted cells progressively adopted zonation profiles matching their new location, with strongly zonated genes showing the greatest adaptive responses. Pharmacological inhibition of EphA2 receptors disrupted zonation, revealing a previously unknown role for epithelial EphA-ephrin-A signaling in regulating enterocyte zonation. These findings demonstrate that self-organization through local epithelial cell-cell communication generates spatial patterns independently of external positional cues or cell-autonomous programs.

## Introduction

The intestinal epithelium exhibits remarkable spatial organization at multiple levels. Most canonical is the organization of different cell types along the crypt-villus axis, with stem cells located at the base of the crypt driving the formation of mature enterocytes that comprise the villus. However, even amongst these villus-localized enterocytes, there is a spatial subpatterning termed zonation, with enterocytes at the villus bottom having different gene expression profiles from those at the villus top. These subpatterns correlate with functional properties: for instance, enterocytes at the villus bottom express antimicrobial peptides, while those at the villus tip express apolipoproteins involved in lipid metabolism. This spatial organization could reflect the distinct challenges faced at different villus positions, with antimicrobial defense prioritized near the crypts to protect stem cells, and lipid processing and secretion enriched at the nutrient-rich villus tip ^1–3^. Yet, while such spatial gene expression patterns have been identified in both mouse and human intestine, it remains unclear whether zonation emerges through epithelial-intrinsic mechanisms or requires continuous instruction from the surrounding tissue.

There are several classes of proposed mechanisms for zone formation. One class is the specification of zone identity by extrinsic signals. For instance, zonated expression profiles could simply be responses to the extracellular environment in the intestine; for example, lipids may be more prevalent near the villus tip, and hence enterocytes in that region respond to those lipids, creating the appearance of zonation. Extrinsic specification could also arise from the orchestration of zonation by morphogen gradients maintained by the mesenchymal tissue ^4,5^. Such morphogen-driven patterning is widespread in development, from neural tube dorsalization to limb bud patterning ^6,7^.

Another class of mechanisms is completely cell autonomous, in which zonation arises from a simple temporal progression of an intrinsic program in a “genetic cascade”. In this scenario, each cell expresses zonation markers in succession. This cascade occurs while the cells move up the villus, and so the apparent zonation profiles would merely reflect the migration of these cells toward the top of the villus during their life cycle. This model, analogous to temporal patterning mechanisms observed in somitogenesis, where a segmentation clock drives spatial pattern formation ^8^ and neurogenesis ^9^, would predict that cells maintain their zonation programs regardless of position.

A third category of mechanisms is that of self-organization. Self-organized patterns form due to local interactions between groups of cells: there is no overarching external “controller” dictating the patterns, nor any intrinsic “program” that is blindly executed over time. Rather, self-organized patterns emerge when cells collectively generate patterns through reciprocal signaling and feedback loops. For example, cells might secrete factors that influence their neighbors’ gene expression, while simultaneously responding to signals from those same neighbors. These interactions create a dynamical system in which the pattern emerges from the collective behavior of the tissue, rather than being imposed by external gradients or predetermined by individual cell programs. Discriminating between these classes of mechanisms is challenging *in vivo* because it is difficult to decouple intrinsic and extrinsic factors that influence zonation. In native intestinal tissue, epithelial cells are constantly exposed to signals from multiple sources: mesenchymal cells secrete morphogens, blood vessels create nutrient and oxygen gradients, neural inputs provide regulatory signals, and mechanical forces from peristalsis and tissue architecture influence cell behavior. These factors are deeply interconnected and cannot be selectively eliminated without disrupting the entire tissue, confounding mechanistic conclusions.

Furthermore, even when molecular signals affecting zonation are identified, perturbation experiments often cannot distinguish between extrinsic, intrinsic, and self-organization models. For example, Beumer et al. ^5^ elegantly demonstrated that BMP signaling regulates enterocyte zonation genes, with BMP inhibition shifting cells toward bottom-zone identity and BMP activation promoting top-zone expression. However, these experiments face two key ambiguities in interpretation.

First, because the experiments lacked spatial measurements, it remains unclear whether BMP perturbations fundamentally disrupted the zonation pattern itself or simply shifted all cells uniformly along the zonation axis while preserving relative spatial order. If zonation order was maintained, this would indicate that while BMP influences zonation states, other mechanisms must establish the spatial pattern. Moreover, the preservation of spatial ordering would still be consistent with either a cell-intrinsic temporal program (where BMP modulates an internal developmental clock) or a self-organizing mechanism (where BMP mediates local cell-cell interactions); thus, these experiments cannot distinguish between these possibilities.

Second, even if BMP establishes zonation patterns, these experiments cannot determine whether BMP acts as a positional cue from external sources, such as mesenchymal cells, or mediates local self-organization within the epithelium. These scenarios make different predictions about scale invariance: if zonation depends on fixed external BMP gradients, villi of different lengths should display different zonation patterns: cells at the top of shorter regenerating villi would lack villus-top identity because they never reach positions with appropriate BMP levels. In contrast, if BMP mediates local epithelial self-organization, zonation could scale proportionally with villus length, allowing even short villi to establish complete zonation patterns. Distinguishing between these mechanisms requires combining spatial measurements with spatial manipulation experiments.

Recent advances in monolayer culture systems have enabled researchers to grow intestinal epithelial cells in two-dimensional formats ^10^, providing a powerful system for testing hypotheses about zonation mechanisms. Originating from intestinal organoids, these monolayers maintain cellular diversity while being far more amenable to imaging than three-dimensional structures, where molecular imaging at single-cell resolution is technically challenging. This two-dimensional nature allows for high-resolution spatial transcriptomic profiling using techniques such as sequential fluorescence *in situ* hybridization (seqFISH), which can detect multiple RNA species at single-cell resolution while preserving spatial information ^11–13^. Equally importantly, the geometric accessibility of monolayers enables targeted spatial perturbations, a critical property for testing mechanistic hypotheses about pattern formation, as demonstrated by classical experiments using spatial rearrangements such as Spemann’s organizer ^14^.

Here, we combined high-resolution spatial transcriptomics with precise spatial manipulations to comprehensively interrogate the spatial organization of gene expression in intestinal epithelial monolayers. We found that intestinal epithelial cells spontaneously establish zonation patterns in the complete absence of mesenchymal, neural, or vascular inputs. Moreover, when cells were transplanted into established monolayers, they adopted expression profiles that matched their new location rather than maintaining their original identity. Together, these findings demonstrate that enterocyte zonation emerges through local epithelial feedback mechanisms independent of external architectural or stromal cues. Pharmacological perturbations implicated Ephrin signaling in this self-organization.

## Results

### 2D intestinal organoid “monolayers” show autonomous zonated gene expression

Intestinal organoids offer a powerful model for studying self-organization, providing experimental control and facilitating imaging while preserving the diversity of epithelial cell types and many aspects of spatial structure. We focused on 2D intestinal monolayers, which are more amenable to imaging and spatial perturbations than traditional 3D organoids. To create monolayers, we dissociated 3D organoids into small groups of cells and plated them on surfaces coated with a thin layer of Matrigel, following established protocols ^10,15,16^. Within 7 days of culture in medium that supports differentiation, these cell clusters expanded into flat epithelial sheets spanning several millimeters. In the native small intestine, enterocytes display zonation - functional and transcriptional variation along the crypt-villus axis ^1^. We examined whether monolayers can recapitulate this zonation despite lacking mesenchymal, neural, and vascular inputs. The presence of zonation in this minimal system would demonstrate that spatial patterning can emerge without external specification signals.

To measure spatial gene expression patterns in this simplified system, we used seqFISH to profile 140 genes at single-cell resolution across 57,549 cells. The panel included genes with known zonated expression in vivo and markers of distinct intestinal cell types. Of the 140 genes, 139 were well detected and used for downstream analysis (Figure 1A; see Methods for panel design).

**Figure 1.**
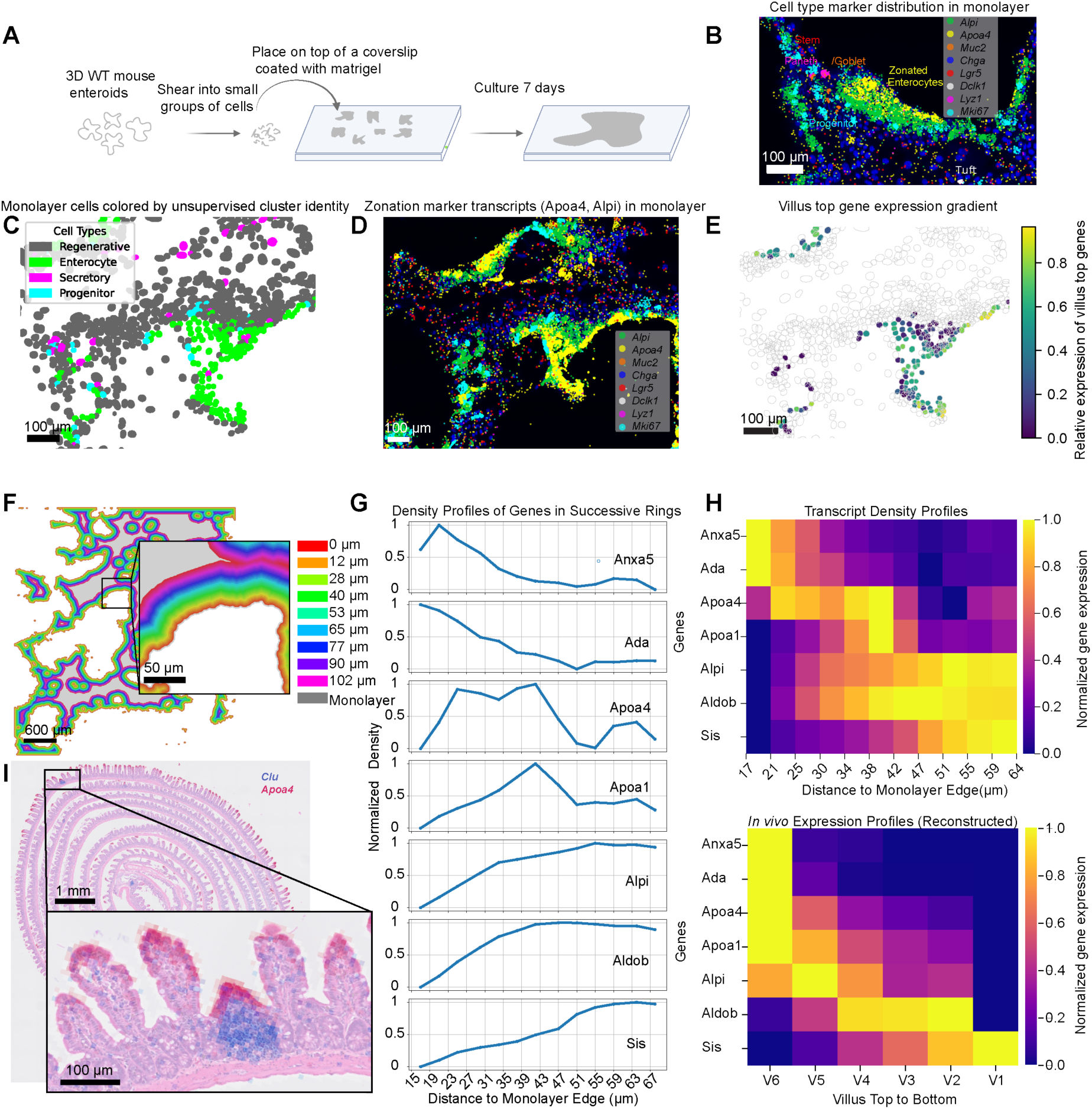
2D intestinal organoids (monolayer) show zonated gene expression. (A) Diagram of experimental procedure. 3D mouse enteroids were mechanically sheared and placed on top of a coverslip coated with a thin layer of Matrigel. Seven days after plating, we recorded and quantified the location of the mRNA expression of a panel of 140 genes using seqFISH. (B) Intestinal monolayer stained with DAPI showing spatial distribution of mRNA molecules for canonical intestinal cell type markers (C) Monolayer cells colored according to cluster identity from unsupervised clustering. (D) Raw monolayer image showing *Alpi* (green) and *Apoa4* (yellow) mRNA expression levels. The scale bar is 100 μm. (E) Villus zonation pattern. Color represents the fraction of top villus marker expression relative to total villus marker expression (top + bottom). Scale ranges from yellow (exclusively top markers) to dark blue (no top markers). (F) Illustration of the computational erosion analysis designed to measure gene expression profiles perpendicular to the monolayer edge. (G) Expression profiles of known zonated genes as a function of distance from the monolayer edge, shown after min-max normalization. (H) Top: normalized expression profiles of selected genes in enteroid monolayers, measured in the direction perpendicular to the monolayer’s edge. Bottom: profiles of the same genes in the small intestine of mice ^1^ along the crypt-villus axis. Both datasets are min-max normalized (scaled to the 0-1 range, where 0 represents the minimum and 1 represents the maximum expression for each gene). (I) Mouse intestine cross-section with spatial expression patterns of *Clu* (blue) and *Apoa4* (red) captured by Visium HD (10X Genomics).

Consistent with previous work ^10^, the monolayers contained all major intestinal epithelial lineages, with cells robustly expressing canonical markers for stem and progenitor cells (*Lgr5*, *Olfm4*), enterocytes (*Alpi*), goblet cells (*Muc2*), enteroendocrine cells (EECs; *Chga*), tuft cells (*Dclk1*), and Paneth cells (*Lyz1*) (Figure 1B and Supplementary Figure 1). These markers were readily detected, and their coordinated expression suggests that the monolayer recapitulates the full spectrum of classical epithelial cell types typically found in the small intestine. Previous reports have highlighted a rare population of regenerative cells ^17,18^ that can emerge in response to tissue damage *in vivo.* Additionally, other studies suggest that organoid cultures can be enriched for cells expressing regenerative signatures ^19^. To measure the presence and quantity of regenerative cell states, we performed unsupervised clustering using gene expression data alone (Figure 1C, see Methods).

This analysis revealed two major transcriptional domains: one resembling normal epithelium, enriched in differentiated and progenitor cell types, and another enriched in regeneration and fetal-associated markers, such as *Clu* and *Msln*. Spatial mapping of these clusters revealed that differentiated enterocytes, goblet cells, and EECs localized to the periphery, while stem cells, progenitor cells, and Paneth cells occupied a medial zone. Cells expressing regeneration markers were concentrated in the central region of the monolayer, consistent with prior observations of injury-like or fetal programs (Figure 1C).

Given that previous reports indicated that zonation depends on external cues ^1,5^, we expected monolayers cultured without any such external cues to show ‘salt and pepper’ expression of *in vivo* zonated genes rather than organized spatial patterns. Surprisingly, intestinal monolayers exhibited zonated expression profiles, despite lacking a full three-dimensional structure and the absence of signaling sources beyond the monolayer itself. Gene expression exhibited a clear spatial organization across the monolayer, with distinct patterns emerging at varying distances from the monolayer edge (Figure 1D-E). For example, *Apoa4* expression (shown in yellow), a top villus marker, was concentrated near the monolayer periphery, while *Alpi* (shown in green), a villus bottom marker, was more prominent in interior regions. This spatial segregation demonstrated that cells adopt different functional states based on their position within the monolayer, reminiscent of the zonation seen along the villus axis *in vivo*. To quantify these patterns, we calculated the ratio of summed expression of top villus genes to the total expression of top and bottom villus genes (Figure 1E). This analysis revealed a clear gradient, with top villus gene expression concentrated at the periphery and bottom villus gene expression increasing toward the center of the monolayer. To reveal spatial organization on a gene-by-gene basis, we defined the distance from the monolayer edge as a zonation axis and systematically measured gene expression profiles inward from the edge (Figure 1F, Methods). Canonical enterocyte genes displayed peak expression in a cascading, layered pattern across the monolayer, mirroring *in vivo* spatial zonation (Figure 1G,H). Among genes in our panel reported to have zonated expression, 73% exhibited patterns consistent with their *in vivo* counterparts. These patterns suggest that the mechanism driving zonation for this subset of genes does not require external signals or a specific three-dimensional geometry.

We sought to determine whether zonation in native intestinal tissue exhibits properties consistent with self-organization. Self-organizing mechanisms can exhibit scale invariance, with patterns adapting proportionally to tissue size through local interactions that establish relative rather than absolute positions ^20,21^. With an extrinsic morphogen gradient mechanism or a cell-intrinsic temporal mechanism, we would expect the villus tip cells of truncated villi not to express the normal villus top signature; rather, we would expect them to be in the same state as cells at an equal distance from the crypt in normal villi. This lack of scaling is due to the fact that both models rely on absolute spatial coordinates, either distance from a morphogen source or migration time from the crypt. In contrast, self-organizing mechanisms in which cells determine their identity through local cell-cell interactions can generate patterns that scale with tissue dimensions, allowing even short villi to establish villus top identities at their tip.

To test whether scaling was observed or not *in vivo*, we analyzed a publicly available Visium HD spatial transcriptomics dataset of mouse small intestine (10x Genomics). We found that the villus length distribution in healthy tissue was relatively homogeneous, making it difficult to test scale invariance under normal conditions. However, we identified several villi that appeared to have undergone injury, as evidenced by their substantially shorter length and high expression of *Clu*, a known marker of regeneration^17,22^ (Figure 1I and Supplementary Figure 4). These regenerating villi were approximately fivefold shorter than adjacent normal villi. Critically, despite their reduced length, these short regenerating villi still established villus-top gene expression states at their tips (Figure 1I and Supplementary Figure 4). The preservation of villus-top identity in regenerating villi of varying lengths supports a self-organizing mechanism in which zonation scales with tissue architecture rather than being determined by fixed positional coordinates.

### Transplanted cells adopt their neighbors’ gene expression patterns

The above results demonstrate that monolayers can exhibit zonation without mesenchymal, neural, or vascular inputs, indicating that zonation can arise independently of externally generated signaling gradients. We next asked whether zonation patterns reflected purely intrinsic programs that execute in a stepwise fashion within individual cells, or if they are a collective property that emerges from local, self-organizing interactions between these epithelial cells. In a cell-intrinsic model, each cell would follow its own internal developmental timer - for example, a newly formed cell at the crypt might be programmed to first express bottom-villus genes, such as Alpi, then gradually switch to expressing top-villus genes, like *Apoa4,* over a fixed time period, regardless of its location. Alternatively, in a self-organizing model, cells would continuously send and receive signals from their neighbors to determine their zonation expression signature. In this scenario, a cell’s zonation identity would depend on its local environment. Observational data alone cannot distinguish between these possibilities; thus, we sought an experimental perturbation that would reposition cells and assess their ability to adapt their zonation identity. As a proxy for such a perturbation, we introduced cells with varied cell types and zonation states into new spatial contexts by dissociating nuclear-GFP-labeled 3D enteroids into single cells and randomly re-seeding them onto pre-formed, differentiated monolayers (Figure 2A). These re-seeded cells then integrate into the monolayer, effectively being “transplanted” into a new location.

**Figure 2:**
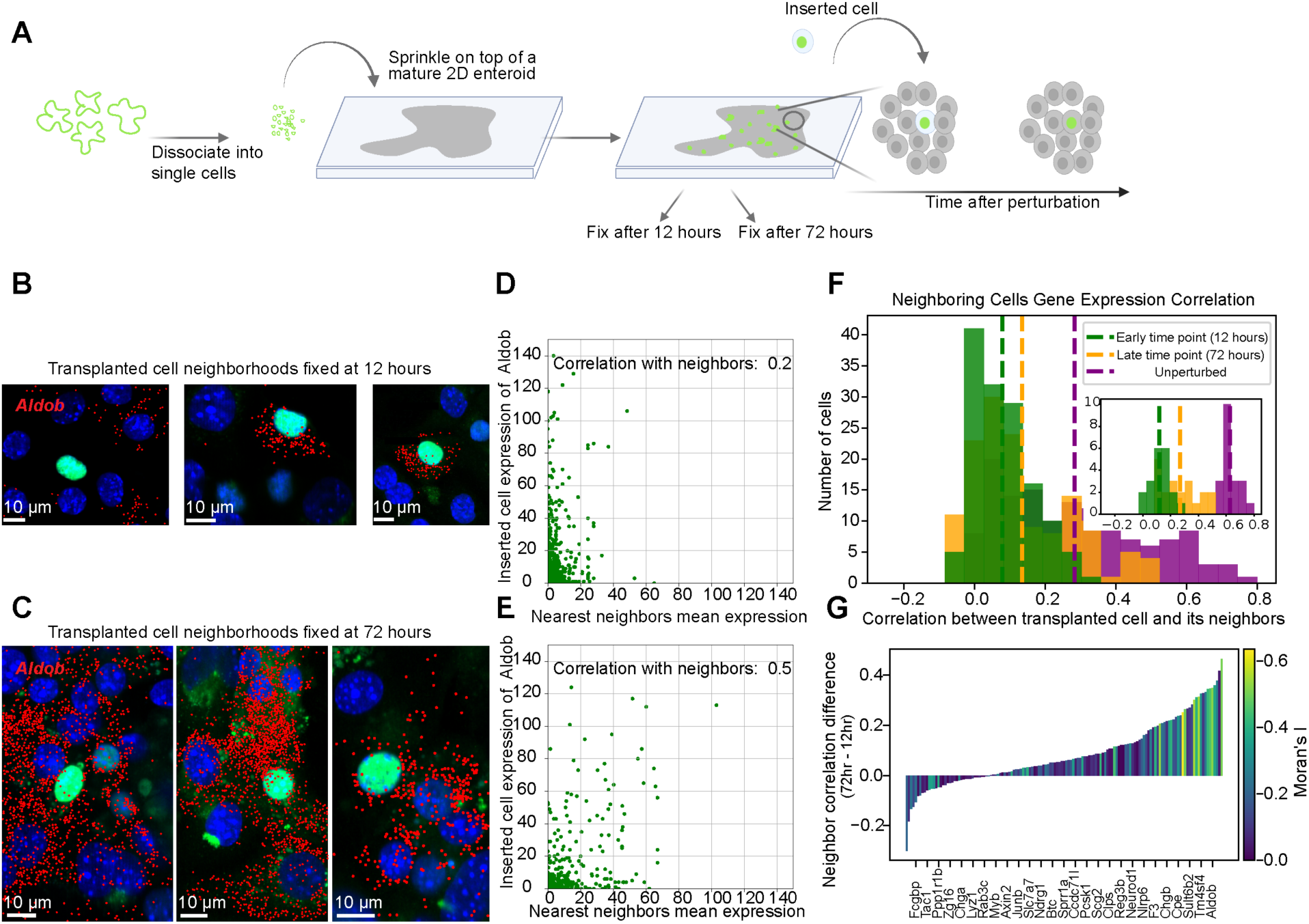
Cells transplanted at paradoxical locations across the monolayer pick up the local gene expression profile. (A) A diagram of the cell transplantation experiment workflow: 3D GFP-labeled organoids are dissociated into single cells and placed on top of a mature intestinal monolayer. GFP-labeled cells integrate into the monolayer at random locations, and the sample is fixed at different time points. (B) Image of a monolayer fixed 12 hours after cell transplantation perturbation. Cell nuclei were stained with DAPI (blue). Transplanted cell nuclei are marked with a green dot, neighboring monolayer cells adjacent to transplanted cells are marked with a magenta dot, and *Aldob* expression is shown in red. (C) Image of a monolayer that was fixed 72 hours after the cell transplantation perturbation (D) Scatter of the *Aldob* count in transplanted cells (y-axis) and the mean count of their neighbors (x-axis) for the 12-hour time point. (E) Scatter of the *Aldob* count in transplanted cells (y-axis) and the mean count of their neighbors (x-axis) for the 72-hour time point. (F) Distribution of the correlation in gene expression of transplanted cells and their neighboring cells over the genes in the panel at 12 hours and 72 hours after the cell transplantation perturbation, compared to the gene expression correlation in the unperturbed monolayer. (G) The difference in correlation between the 72-hour and the 12-hour time point, per gene. Bars are colored by the gene’s Moran’s I value.

We hypothesized that this spatial perturbation could lead to one of two outcomes. In a **cell-intrinsic** mechanism, transplanted cells would retain their transcriptional profiles, independent of their new environment, and potentially change over time according to internal programs. In contrast, a **local feedback** mechanism predicts that transplanted cells would be affected by their surroundings, for example, by adapting their gene expression to more closely match the zonation profile of the surrounding tissue over time. We saw evidence for the latter scenario. For instance, representative examples of *Aldob* expression in neighborhoods around transplanted cells at early and late time points are shown in Figures 2B and 2C. Qualitatively, the expression levels of the transplanted cells increasingly matched those of their neighbors over time. Quantitatively, the correlation between *Aldob* expression in transplanted cells and their immediate neighbors increased from 0.2 at early time points (Figure 2D) to 0.5 at later time points (Figure 2E). Extending this analysis across the full 140-gene panel, we observed a significant global increase in correlation values over time (Figure 2F, *p* < 1e−6), consistent with a plasticity-driven mechanism of zonation.

Note that an alternative interpretation is that transplanted cells may preferentially incorporate into specific zones, artificially inflating their similarity to neighboring cells. However, under such a selection mechanism, gene expression correlations between transplanted cells and their neighbors would remain stable over time. In contrast, a plasticity mechanism would predict increasing correlation with time, which is far more consistent with our observations over many genes.

Calculating the difference in correlation values between time points, we found that for most genes in our panel, the difference was positive (73%, Figure 2G), indicating that the correlation is higher at the later time point. We next asked whether genes showing the strongest adaptive responses shared common characteristics. Given our evidence that zonation is driven by local cell-cell communication, we hypothesized that genes with the largest increases in correlation, i.e., those most responsive to their new neighborhoods, would exhibit strong zonation patterns. To test this hypothesis, we ranked genes based on their spatial autocorrelation in the monolayer using Moran’s I, which measures the degree to which gene expression is spatially ordered rather than randomly distributed. Genes with zonated spatial expression patterns exhibited high Moran’s I scores, whereas those with random or uniform expression patterns showed low scores. Consistent with our hypothesis, we found that genes with high Moran’s I values tended to exhibit the largest changes in correlation coefficient (Figure 2G), indicating that zonated genes were particularly influenced by local interactions.

### Transplanted cells adopt the zonation profile of their neighbors

In the previous section, we established that transplanted cells become more similar to their neighbors over time, as indicated by comparisons of the expression of individual genes. This plasticity supports a self-organization mechanism over cell-intrinsic temporal programs. However, while transplanted cells became increasingly similar to their neighbors in terms of global correlation over gene expression (reflecting an averaged measure), a complete test of the self-organization mechanism requires examining the extent of similarity, specifically of zonation programs, between transplanted and neighboring cells. To assess whether transplanted cells adopt coordinated zonation patterns that matched their neighbors, we first characterized the spatial zones in unperturbed monolayers. We used the gene expression profiles binned by distance from the monolayer edge to define discrete “zones” analogous to positions along the crypt-villus axis (Figure 1E,F). This approach generated a spatial zone atlas based on the expression patterns of canonical enterocyte genes. As shown in Figure 1E-F, this atlas revealed that top villus genes are expressed closer to the monolayer edge, while bottom villus genes are expressed further inward (Figure 1D), patterns that closely match *in vivo* zonation along the villus axis ^10^, Figure. 1H).

We then projected the profiles of both transplanted and neighboring cells onto this atlas, resulting in a zone probability for each cell (see Methods, Figure 3A). To verify this mapping, we first applied it to unperturbed monolayers, finding that the expected zone identity of each cell also followed a spatial profile indicative of ordered zonation, as observed *in vivo* (Figure 3B).

**Figure 3:**
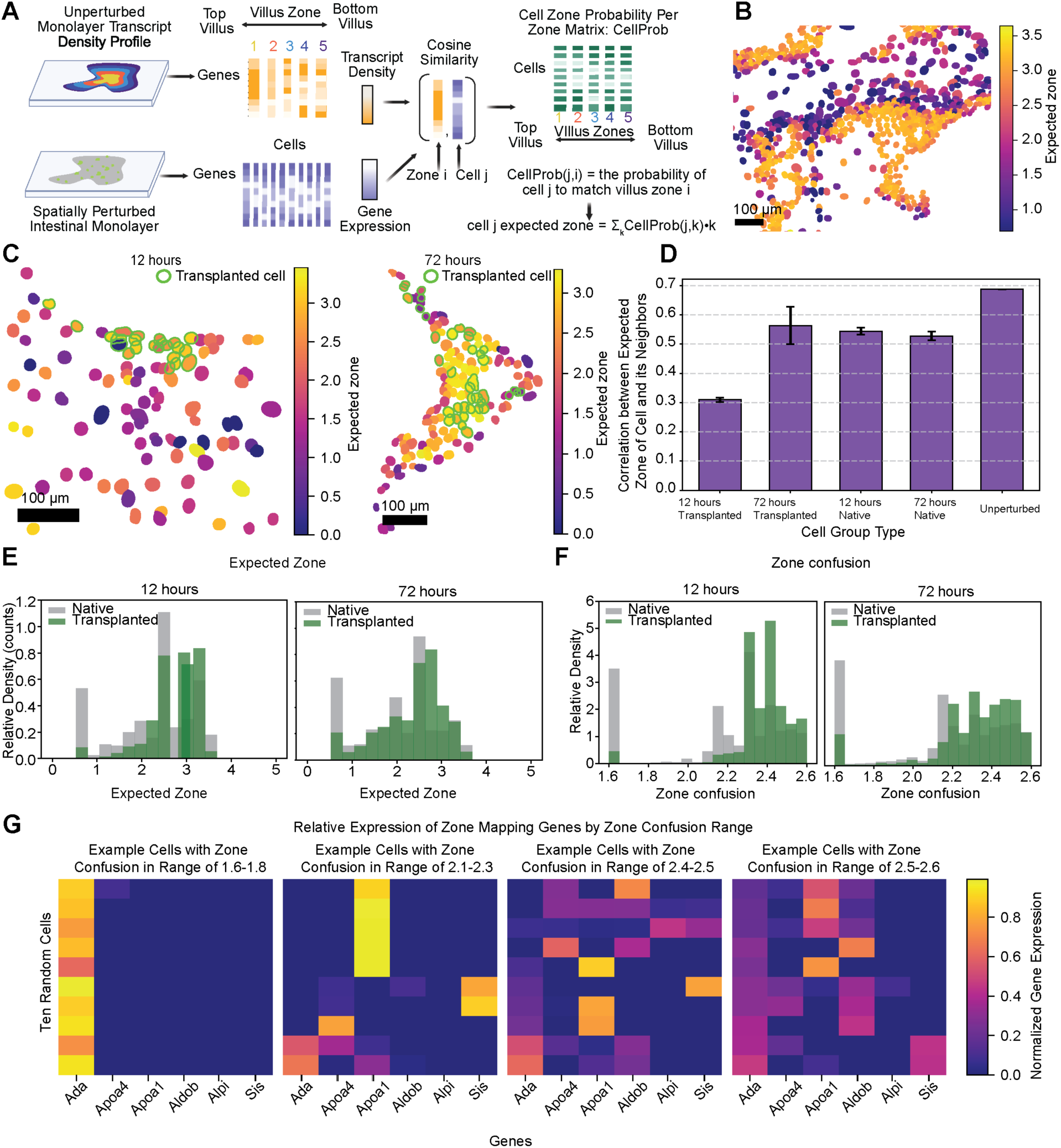
Transplanted cells adopt the zonation profile of their neighbors. (A) Schematic diagram of expected zone calculation. Gene expression profiles from unperturbed monolayers are binned by distance from the edge to create a spatial zone atlas. For each cell in perturbed monolayers, cosine similarity is calculated between the cell’s expression profile and each zone’s profile, yielding a probability distribution over zones. The expected zone is the weighted average of zone identities, where higher values indicate top villus identity and lower values indicate bottom villus identity. (B) Representative unperturbed monolayer segment with cells colored by their expected zone value. Color scale ranges from 0.0 (bottom villus, purple) to 3.5 (top villus, yellow). (C) Spatial distribution of expected zone values in transplanted monolayers at 12 hours (left) and 72 hours (right) post-transplantation. Transplanted cells are circled in green. At 12 hours, transplanted cells (predominantly yellow/orange, zones 2.5-3.5) show poor correspondence with their local neighborhoods. (D) Quantification of expected zone correlations across conditions. Bar height represents the Pearson correlation coefficient between cells and their local neighbors; error bars show the standard error. Transplanted cells show lower correlation with neighbors at 12 hours. (E) Distribution of expected zone values. Histograms show the relative density of cells assigned to each zone (0-5 scale) for transplanted cells (green) versus resident monolayer cells (gray) at 12 hours (left) and 72 hours (right). (F) Zone confusion distributions. Zone confusion is defined as the entropy of each cell’s zone probability distribution: H = -Σ P(zone_i) × log₂(P(zone_i)). Higher values indicate cells with mixed/ambiguous zone identity (high entropy, expressing genes from multiple zones), while lower values indicate cells with clear commitment to a single zone (low entropy). Histograms compare transplanted cells (green) to resident cells (gray) at 12 hours (left) and 72 hours (right). (G) Gene expression patterns underlying zone confusion. Heatmaps show normalized expression of the six zone-mapping genes (*Ada, Apoa4, Apoa1, Aldob, Alpi, Sis*) in example cells binned by their zone confusion values. Genes are ordered by their typical expression along the villus axis (*Ada* at top/edge, *Sis* at bottom/center). Each column represents one cell; expression values are normalized per cell (sum to 1) to show relative contributions of each gene.

We then analyzed the expected zones of transplanted cells compared to their neighboring cells. Figure 3C shows representative zone mapping of monolayer subsections at 12 hours (left) and 72 hours (right) post-transplantation. At the early time point (12 hours), the transplanted cells predominantly mapped to zone 3 (yellow and orange), while their neighbors occupied the lower zones (pink and purple) in this region. By 72 hours, the transplanted cells showed markedly greater similarity to their neighbors’ zones. Quantifying this effect, we found that the correlation between the expected zone values of transplanted cells and their neighbors increased from R = 0.31 at 12 hours to R = 0.56 at 72 hours (both p < 0.001; Figure 3D).

In addition to the increasing correlation values over time, we observed that the transplanted cells occupied a wider range of zone identities at 72 hours compared to 12 hours. This expansion in zone diversity further supports adaptation of transplanted cells to match the diverse zones of their local neighborhoods across the monolayer (Figure 3E).

Having established that transplanted cells adopt the zone signatures of their neighbors, we next asked whether transplanted cells achieve zone identities as distinct as those of resident monolayer cells. To address this, we defined a measure of “zone confusion” based on the entropy of the zone distribution, providing a holistic measure of how consistently cells express zone-specific gene programs (Figure 3F). Cells with low zone confusion expressed genes predominantly from a single zone (expressing few zone-mapping genes at high levels), while cells with high zone confusion co-expressed genes from multiple zones simultaneously, creating mixed expression profiles (Figure 3G). Cells with low zone confusion expressed genes predominantly from either the bottom, middle, or top villus zones, while cells with high zone confusion co-expressed genes from multiple zones, creating mixed expression profiles (Figure 3F).

We found that the extent of zone confusion in transplanted cells in both 12-hour and 72-hour monolayer was significantly higher than in non-transplanted monolayer cells (12 hours: transplanted cells mean entropy = 2.4 ± 0.2, n = 2,879; native cells mean entropy = 2.2 ± 0.3, n = 13,108; KS test, p < 1×10⁻²⁴⁰; 72 hours: transplanted cells mean entropy = 2.3 ± 0.2, n = 1,054; native cells mean entropy = 2.16 ± 0.32, n = 13,104; p < 1×10⁻⁶⁰). This higher zone confusion indicates that while transplanted cells are adapting to the profiles of their neighbors, that adaptation is only partial, perhaps reflecting transient intermediate states. Supporting the notion that it is a transient state, we found that the difference in confusion between transplanted and non-transplanted cells was smaller in the 72-hour transplanted cells than 12-hour transplanted cells (Figure 3F).

### Pharmacological perturbations uncover the molecular pathway driving zonation

Having demonstrated that zonation emerges without externally-specified signaling gradients and that transplanted cells adopt local expression patterns rather than following intrinsic programs, we sought to identify the molecular pathways that mediate this self-organization. We analyzed receptor-ligand pairs within the enterocyte cluster using CellPhoneDB ^23^ on a single-cell RNA-seq dataset from intestinal monolayers (see Methods). This analysis yielded 22 significant receptor-ligand interactions (see supplementary table 1 for the full interaction list). From these interactions, we prioritized pathways with established roles in pattern formation. BMP and Ephrin emerged as top candidates (see Methods for more details), specifically BMP2-BMPR1A, EphA2-Efna1, and EphA1-Efna1. We expanded our candidate list to include WNT, a major signaling pathway in both intestine and liver that was recently shown to drive liver zonation ^24^, and Notch, which mediates self-organized pattern formation in other intestinal contexts ^25^. BMP was a strong candidate as it has already been linked to intestinal zonation ^5^. The Ephrin pathway is particularly intriguing. While EphB2 and EphB3 interact with ephrin-B ligands in the intestinal crypt to mediate cell sorting and prevent intermingling between proliferative and differentiated cells ^26^, the role of Ephrin-A receptors and their ligands in the intestine remains unclear.

To test whether any of these pathways are required for zonation, we cultured monolayers for seven days with pathway-specific antagonists: LDN-193189 (BMP inhibitor), IWP-2 (Wnt inhibitor), ALW-II-41-27 (Ephrin receptor inhibitor, selectively targets EphA2), and RO4929097 (Notch inhibitor). We then fixed the monolayers and profiled *Sis* (villus bottom) and *Apoa4* (villus top) expression using HCR-FISH, followed by computational erosion analysis to measure expression profiles perpendicular to the monolayer edge.

Monolayers treated with LDN-193189 (BMP inhibitor), IWP-2 (Wnt inhibitor), or RO4929097 (Notch inhibitor) showed zonation patterns qualitatively similar to control (ENR) media: edge cells expressed *Apoa4* but not *Sis*, while interior cells expressed *Sis* but not *Apoa4* (see Figure 4 A and B and Supplementary Figure 2). In striking contrast, ALW-II-41-27 (Ephrin inhibitor) treatment produced dramatically altered patterns (Figure 4 C and D). Ephrin inhibitor treatment resulted in a marked increase in *Sis*-expressing cells (villus bottom marker), both in absolute number and relative to *Apoa4*-positive cells (villus top marker), compared to all other conditions, while *Msln*-expressing cells (regenerative marker) remained unchanged (Figure 4E). Additionally, edge cells co-expressed both top and bottom villus markers within individual cells, thereby disrupting the mutual exclusivity of these markers and the spatial segregation that normally restricts them to distinct zones (Figure 4G).

**Figure 4:**
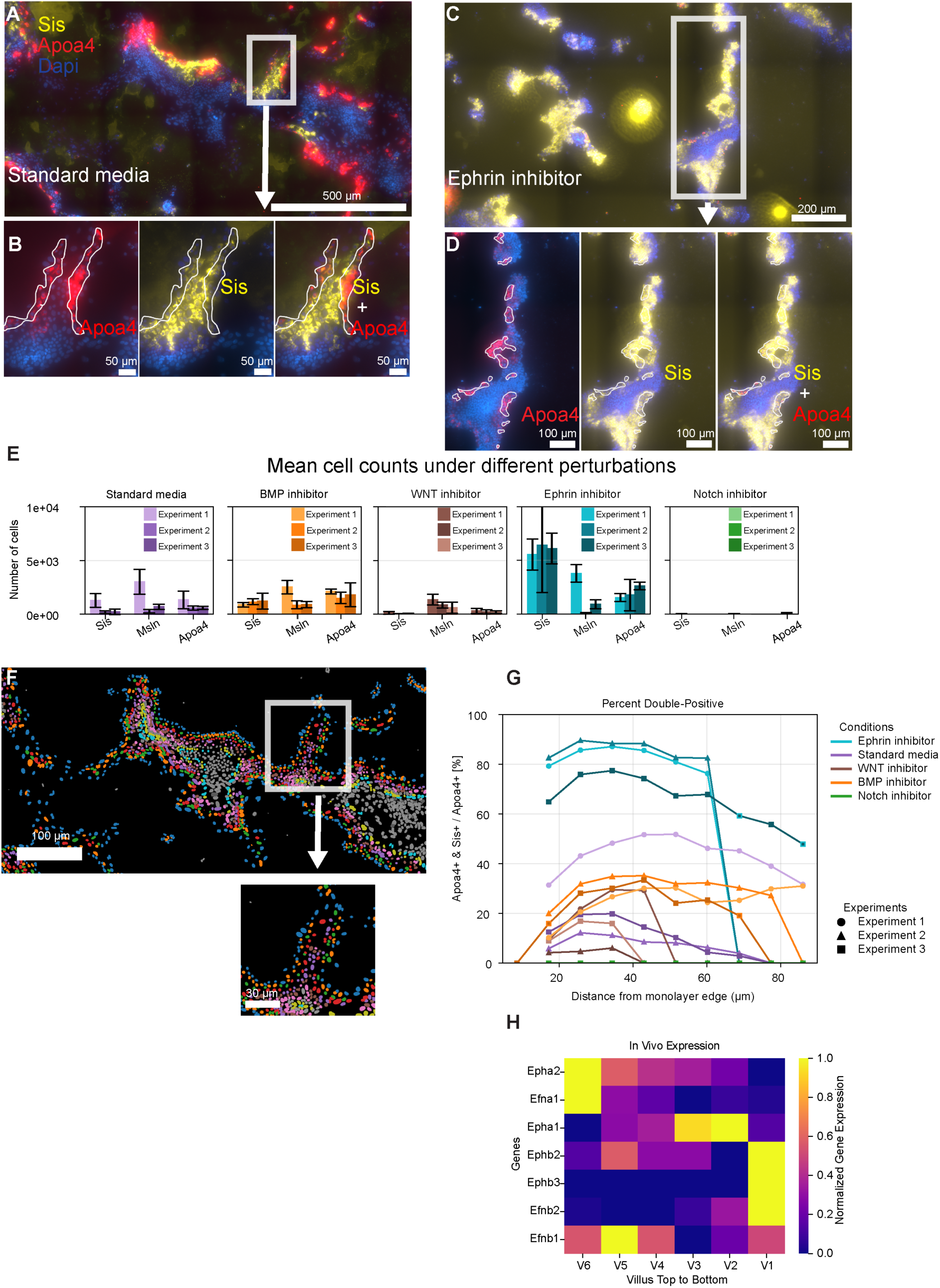
Pharmacological perturbation shows EphA is required for proper zonation. (A) Monolayer under normal conditions. *Sis* (yellow) is a villus bottom gene, *Apoa4* (red) is a villus top gene. (B) zoomed-in images of A. The area of *Apoa4* high expression is circled in white. (C) Monolayer under Ephrin inhibitor. (D) zoomed-in images of (C). (E) Mean number of cells positive for *Sis*, *Msln* (regenerative marker), and *Apoa4* per monolayer. Cells were scored as positive if fluorescence intensity exceeded background + 2 standard deviations. Each bar color represents an independent biological replicate, with n = 4 monolayers per replicate (n = 3 biological replicates). Error bars indicate standard deviation across monolayers within each replicate. (F) Computational erosion analysis: cells are labeled according to their distance from the edge. (G) Percentage of cells positive for both *Apoa4* and *Sis* out of *Apoa4* positive cells along spatial bins created by the computational erosion analysis (H) *In vivo* expression of the Ephrin pathway receptors and ligands in the small intestine of mice, data taken from ^1^

Given the pronounced effect of ALW-II-41-27 on zonation, we examined the spatial expression pattern of the EphA2-EfnA1 receptor-ligand pair along the crypt-villus axis *in vivo*. Using the published RNA-seq dataset from ^1^, we found that both EphA2 and EfnA1 are predominantly expressed at the villus tip, precisely the region corresponding to monolayer edge cells where Ephrin inhibition caused aberrant co-expression of top and bottom villus markers (Figure 4H). This spatial correspondence between *in vivo* EphA2-EfnA1 expression and the site of disrupted zonation in our perturbation experiments supports a direct role for this signaling axis in maintaining villus tip identity.

### The regenerative response is spatially continuous

The intestine has extensive regenerative capabilities. In recent years, several studies using various perturbations (irradiation, chemical agents, pathogens) have uncovered intestinal cell populations expressing “fetal” or “regenerative” transcriptional signatures that emerge in response to these perturbations and are linked to regeneration ^17,18,22^. It is hypothesized that intestinal regeneration requires cells to transition through a regenerative state. However, it remains unclear whether this regenerative gene expression signature marks a distinct cell population or represents a cellular response to injury (Figure 5A).

**Figure 5:**
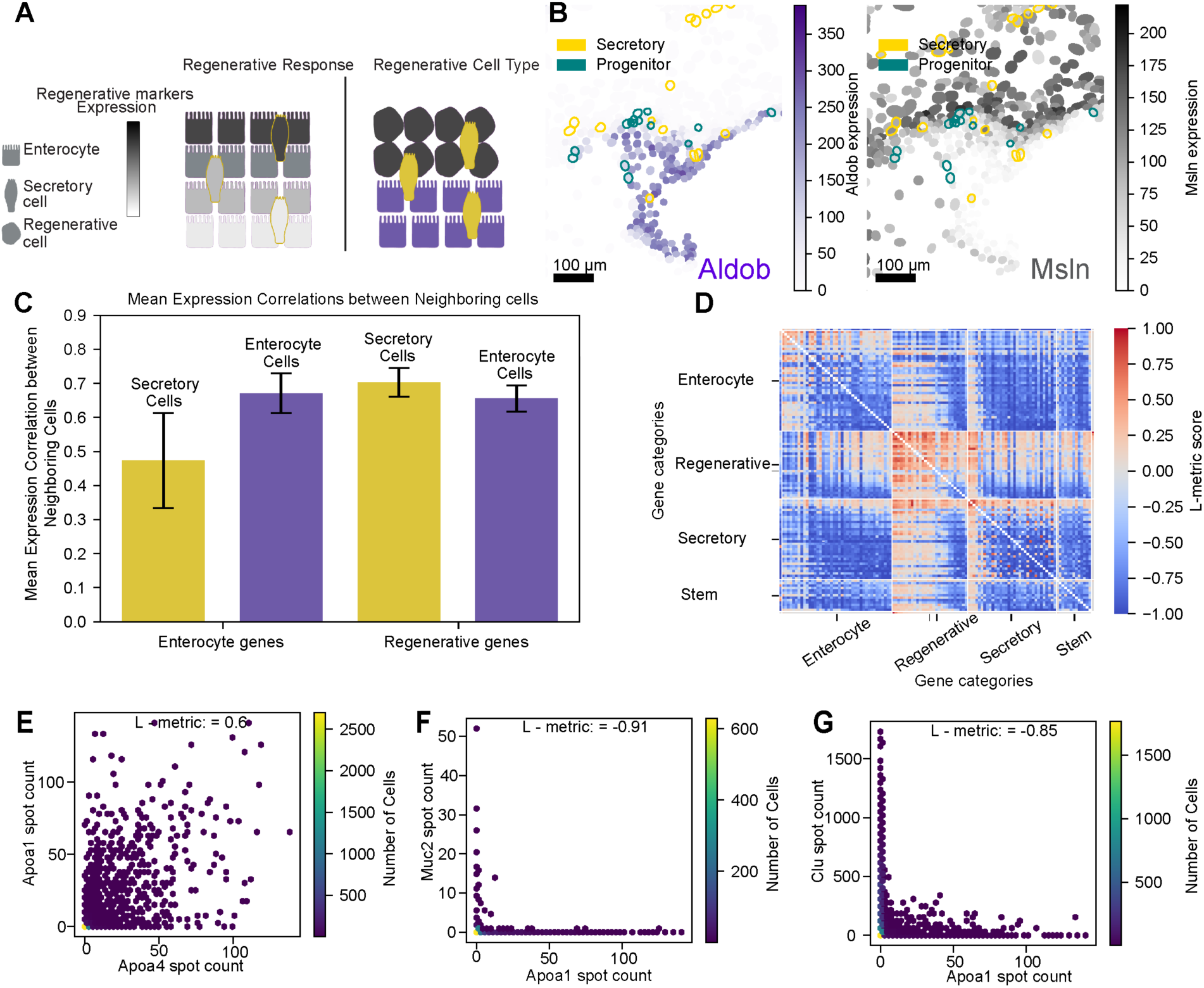
The regenerative response is spatially continuous. (A) The two hypothetical spatial arrangements of “regenerative” and “normal” epithelial cells in the monolayer. (B) Right: Spatial expression of *Aldob*, which is usually expressed in enterocytes; Left: Spatial expression of *Msln*, marking a regenerative state. (C) The regenerative state is smooth, regardless of cell type: gene expression correlates with the local neighborhood of either enterocyte or secretory cell types. (D) Pairwise L-metric values for the full 140-gene panel. Genes are arranged into four clusters: enterocyte, regenerative, secretory, and stem. (E) - (G) Examples of three archetype L-metric patterns.all cell types.

We observed that certain areas within the intestinal monolayer display regenerative characteristics (Figure 1C) and express genes considered part of the regenerative signature, including *Clu*, *Msln*, and *Basp1* ^17^. A key question is whether these regenerative markers define a discrete cell population or represent a transient cellular state. If regenerative cells were a distinct cell type, we would expect sharp boundaries between regenerative and normal epithelial regions, similar to the clear demarcation between enterocytes and goblet cells (illustrated in Figure 5A). Instead, we observed smooth, gradual transitions in marker gene expression between regenerative and normal regions (Figure 5B). For example, *Msln* (regenerative marker) and *Aldob* (enterocyte marker) show overlapping expression in intermediate zones rather than mutually exclusive domains. This continuity in cellular state suggests that regenerative gene expression represents a cellular response state that cells can tune, rather than a fixed cell-type identity.

To quantify this observation, we employed two analytical approaches. First, we assessed the spatial continuity of gene expression by computing correlations between adjacent cells within neighborhoods containing both enterocytes and secretory cells. This approach leverages the fact that secretory cells, even when embedded within enterocyte populations, should not express enterocyte-specific markers if these markers truly define cell identity. Conversely, we hypothesized that regenerative genes would exhibit spatially smooth expression patterns across all cell types.

Our analysis confirmed this hypothesis: regenerative markers maintained high correlation values between adjacent cells regardless of cell type (enterocyte or secretory), indicating continuous spatial expression. In contrast, enterocyte markers showed minimal correlation in enterocyte-secretory cell pairs, consistent with their role as cell-type-specific markers (Figure 5C).

Second, we developed a quantitative measure termed the “L-metric” (^27^, see Methods) to evaluate the degree of mixed expression for gene pairs, where the underlying assumption is that genes acting as true cell-type markers should not be co-expressed with markers of other cell types in fully differentiated cells. As shown in Figure 5D, quantifying the L-metric between all gene pairs organized by cell-type clusters reveals that regenerative marker genes frequently co-express with cell-type markers across various cell populations within the monolayer, whereas cell-type marker genes rarely co-express with each other. This distinctive pattern suggests that the regenerative state differs fundamentally from discrete cell types, exhibiting continuous expression patterns across diverse cell populations. Furthermore, regenerative gene expression demonstrates spatial continuity throughout the monolayer.

## Discussion

In this study, we sought to distinguish between three possible mechanisms for intestinal zonation: extrinsic specification by environmental signals, cell-intrinsic temporal programs, or self-organization through local cellular interactions. We demonstrate that intestinal epithelial cells can establish zonation patterns in 2D monolayers without requiring mesenchymal, neural, or vascular inputs, indicating that zonation can arise independently of these extrinsic factors. Using spatial transcriptomics, we found that 73% of zonated genes exhibited expression patterns consistent with their *in vivo* patterning, despite the absence of external architectural cues and different geometry compared to native tissue. When cells were transplanted into established monolayers, they progressively adopted zonated gene expression profiles that matched their new location, ruling out cell-intrinsic temporal programming as the primary mechanism. If cells followed a predetermined temporal program analogous to those in somitogenesis or neurogenesis, they would be expected to maintain their expression patterns regardless of position. Instead, the correlation between transplanted cells and their neighbors increased for most genes over time post-transplantation, with zonated enterocyte genes showing the strongest adaptive responses. These findings establish that intestinal zonation in monolayer organoids emerges through self-organization, where local epithelial feedback mechanisms generate spatial patterns without requiring external tissue architecture or predetermined cellular programs. While extrinsic signals may modulate zonation in vivo, our results demonstrate that self-organizing mechanisms within the epithelium are sufficient to establish zonation patterns. It is possible that self-organization provides a layer of redundancy with external specification in the natural context.

Further supporting the self-organization model, zonation patterns in the mouse intestine appear to exhibit scale invariance, a hallmark of self-organized systems. Regenerating villi, identified by *Clu* expression, are substantially shorter and less ordered than normal villi, yet they still establish villus top gene expression states at their tips (See Supplementary Figure 4). If zonation were driven by external gradients emanating from the crypt base or by cell-intrinsic temporal programs linked to migration distance, we would expect cells at the top of these shortened regenerative villi to express the same markers as cells at equivalent heights in neighboring normal villi, essentially displaying mid-villus rather than villus-tip identity. Instead, the preservation of tip identity despite altered villus geometry suggests that zonation scales with the available tissue architecture. This scale invariance is consistent with self-organizing systems where local interactions establish relative positional identities independent of absolute spatial coordinates, similar to how reaction-diffusion systems can generate proportionally scaled patterns across different tissue sizes

Our pharmacological perturbations revealed that Ephrin signaling plays a critical role in maintaining zonation patterns. The Eph-ephrin pathway is known to drive cell sorting in the intestine, with EphB2 and EphB3 receptors (highly expressed in crypt cells) and ephrin-B ligands (highly expressed in villus cells) exhibiting complementary expression that prevents intermingling between proliferative and differentiated compartments ^26^. Our results suggest that Ephrin signaling also regulates zonation within the villus compartment, but through a distinct mechanism involving EphA2 receptors and ephrin-A ligands. Analysis of published *in vivo* RNA-seq data ^1^ revealed that EphA2 and multiple ephrin-A ligands, including Efna1, are co-expressed at the villus tip, in contrast to the complementary expression pattern of EphB2/B3-ephrin-B pairs along the crypt-villus axis. This co-localization suggests that EphA2-ephrin-A signaling may underpin mechanisms mediating behaviors other than cell sorting.

If zonation were achieved purely through Ephrin-mediated segregation of pre-specified cell types, we would expect EphA2 inhibition to produce a “salt and pepper” distribution of fully-committed top or bottom cells. Instead, we observed individual cells co-expressing both top and bottom villus markers at the monolayer edge, indicating that cells lost their distinct zonation identities rather than being mis-sorted. This phenotype, which occurs specifically at edge locations where EphA2 and ephrin-A ligands are normally co-expressed *in vivo*, suggests that EphA2-ephrin-A signaling directly influences gene expression programs in a position-dependent manner, rather than merely preventing cell intermingling.

One possibility is that co-localized EphA2-ephrin-A signaling at the villus tip/monolayer edge functions as a boundary-sensing mechanism, with cells requiring local EphA2 activation to maintain top-zone identity. This result would be consistent with the known role of Eph-ephrin signaling in establishing tissue boundaries, but represents a conceptually different mode of action: rather than preventing mixing between two pre-determined populations (as EphB2/B3-ephrin-B does in the crypt-villus axis), EphA2-ephrin-A signaling may actively instruct and maintain villus tip gene expression programs through local cell-cell communication. However, further mechanistic work is needed to determine which ephrin-A ligand(s) interact with EphA2 in this context, whether EphA2-ephrin-A signaling acts through contact-dependent forward and/or reverse signaling, whether it responds to edge geometry or cell density, and how it interfaces with other zonation regulators such as BMP signaling.

The ability of monolayers to recapitulate in vivo zonation patterns despite their different geometry, a two-dimensional flat, expanding disc rather than the finger-like 3D projections of intestinal villi, suggests that zonation mechanisms can operate across diverse tissue architectures. Previous work has identified zonation patterns in both mouse and human intestine ^1,2^, but our findings indicate that the epithelium possesses the capacity for spatial self-organization that is independent of external positional cues or tissue architecture. However, the zonation patterns form at the edge of the epithelial cell monolayer, suggesting that zonation is affected by local geometry.

Our analytical approaches, including the expected zone calculation and L-metric, provide quantitative frameworks for analyzing spatial patterns of gene expression. The expected zone metric allowed us to assign cells probabilistically to positions along a one-dimensional zonation axis, capturing not only mean positions but also the confidence of zone assignments through entropy calculations. Cells with high zone confusion (entropy) often represented transitional states, providing insights into the dynamics of spatial reprogramming. These methods complement existing spatial analysis tools ^11,13^ and can be applied to study spatial organization in other tissues.

Our analysis reveals that the regenerative state in intestinal epithelium represents a continuous spectrum rather than a discrete cell type. Cells expressing regenerative markers such as *Clu*, *Msln*, and *Basp1* ^17^ showed smooth spatial transitions and frequently co-expressed with markers of differentiated cell types, contrasting with the traditional view of regenerative cells as a distinct population ^22^. While enterocyte markers rarely co-express with secretory cell markers, regenerative markers readily co-express with multiple cell-type markers across the monolayer, suggesting that regeneration represents a cellular state rather than a fixed identity. This continuous nature implies that the intestine could leverage the plasticity of existing cells to mount regenerative responses through local interactions, potentially allowing tissue-wide responses to injury without requiring pre-existing specialized cells. This view is consistent with recent observations ^18,28^ of injury-induced plasticity in the intestine, but it provides an alternative framework for understanding how regenerative programs may spread through spatial continuity.

Our monolayer model, while revealing self-organizing principles of zonation, does not capture all factors that may influence spatial patterning *in vivo*. The absence of blood vessels eliminates oxygen and nutrient gradients that can contribute to zonation ^4^, and the lack of neural inputs removes additional potential regulatory mechanisms. A key future direction will be to understand how these extrinsic factors interact with the self-organizing properties we have identified, whether they modulate, enhance, or override local epithelial feedback. Additionally, our observations were limited to 72 hours post-transplantation, leaving questions about the long-term stability of adopted cell identities and whether cells can undergo multiple rounds of reprogramming. Interestingly, the capacity for self-organized zonation may vary across tissues; for example, liver organoids appear to lose the characteristic metabolic zonation observed *in vivo* ^29–31^, suggesting that hepatocyte zonation may be more dependent on extrinsic signals than intestinal zonation. Similar self-organizing principles may apply to other zonated tissues more broadly.

### Experimental Methods

#### Small Intestinal Crypt Isolation

Mouse small intestinal crypts were isolated using a modified protocol adapted from previous methods. Briefly, mice were euthanized, and the small intestine was dissected from the pyloric sphincter. The proximal 10 cm of small intestine was harvested, removing all associated fat, connective tissue, and vasculature. The tissue was immediately placed in ice-cold wash buffer (HBSS supplemented with 50 μg/mL gentamicin).

The intestinal segment was cut to 5 cm, opened longitudinally, and the luminal surface was gently scraped with a glass coverslip under a dissecting microscope to remove villi. The tissue was then transferred to a 50 mL conical tube containing wash buffer and pipetted 5 times using a serological pipette. This washing step was repeated until the supernatant was clear (typically 3 times).

For crypt dissociation, the tissue was incubated in dissociation buffer (wash buffer supplemented with 5 mM EDTA) on a shaking platform at 4°C for 15 minutes. Following incubation, the tissue was gently pipetted 20 times, and the supernatant was discarded. Crypts were mechanically isolated by resuspending the tissue in 10 mL isolation buffer (wash buffer with 0.5% BSA) and vigorously pipetting 20 times with the serological pipette held at the bottom of the tube. The crypt-containing supernatant was filtered through a 70 μm cell strainer, and crypts were counted using an inverted microscope. Crypts were pelleted by centrifugation at 300g for 5 minutes.

### Organoid Culture Establishment

For initial plating, 50-200 crypts were resuspended per 25 μL of growth factor-reduced Matrigel (Corning) and plated as domes in the center of 8-well chamber slides (Ibidi). The Matrigel was polymerized at 37°C for 30 minutes before adding 200 μL of culture medium.

Mouse small intestinal organoids were maintained in either housemade medium or IntestiCult™ Organoid Growth Medium (STEMCELL Technologies). Housemade medium consisted of Advanced DMEM/F12 supplemented with 1× B27 (Gibco), 1× N2 (Gibco), 1× GlutaMAX (Gibco), 10 mM HEPES, 1 mM N-acetylcysteine, 50 ng/mL murine EGF (PeproTech), 100 ng/mL murine Noggin (PeproTech), and 500 ng/mL murine R-spondin 1 (R&D Systems). Both media were supplemented with 1× penicillin-streptomycin and gentamicin. For the first 48 hours of culture, the medium was supplemented with 10 μM Y-27632 (a ROCK inhibitor, Selleck Chemicals). Medium was replaced every other day.

### Organoid Passaging

Organoids were passaged every 5-7 days using TrypLE Express (Gibco) following established protocols. Culture medium was removed, and 400 μL of TrypLE Express supplemented with 10 μM Y-27632 was added to each well. The Matrigel domes were mechanically disrupted using a P1000 pipette, and the suspension was transferred to a 15 mL conical tube. For multicellular fragments, samples were incubated at 37°C for 5 minutes. For single-cell dissociation, incubation was extended to 20 minutes. The enzymatic reaction was quenched with 10 mL of cold basal medium. For single-cell preparations, the suspension was filtered through a 40 μm strainer. Cells were pelleted at 400g for 5 minutes and resuspended in fresh Matrigel for replating at a 1:3 to 1:4 split ratio.

### Cryopreservation

For long-term storage, organoids were cryopreserved using Recovery™ Cell Culture Freezing Medium (Gibco). The medium was removed from the culture wells, and 400 μL of ice-cold freezing medium was added. The Matrigel-organoid suspension was gently disrupted and transferred to cryovials (1 well per vial). Vials were stored at -80°C in controlled-rate freezing containers for 3 days before transfer to liquid nitrogen. For thawing, cryovials were rapidly warmed in 37°C water, diluted with 10 mL of cold basal medium, and centrifuged at 300g for 5 minutes before being resuspended in Matrigel.

### Single-Cell RNA Sequencing and Unsupervised Clustering of Intestinal Monolayers

#### Sample Preparation and Library Generation

Two-dimensional intestinal epithelial monolayers were cultured as previously described and harvested at day 2 post-plating at two different seeding densities (high and low). Monolayers were dissociated into single-cell suspensions using TrypLE Express (Gibco) supplemented with 10 μM Y-27632 ROCK inhibitor at 37°C for 20 minutes. The enzymatic reaction was quenched with cold basal medium, and cells were filtered through a 40 μm strainer to ensure single-cell preparations. Cell viability and concentration were assessed using the trypan blue exclusion method.

Single-cell RNA sequencing libraries were prepared using the 10x Genomics Chromium Single Cell 3’ v3.1 platform according to the manufacturer’s protocol. Briefly, single-cell suspensions were loaded onto the Chromium Controller to generate gel bead-in-emulsions, targeting recovery of approximately 10,000 cells per sample. Following reverse transcription and cDNA amplification, libraries were prepared and sequenced on an Illumina NovaSeq 6000 platform with a target sequencing depth of 50,000 reads per cell.

### Data Processing and Quality Control

Raw sequencing data were processed using Cell Ranger v6.0 (10x Genomics) with alignment to the mouse reference genome (mm10). Count matrices for each sample were generated and imported into R (v4.1.0) using Seurat (v4.1.0). Quality control filtering was applied to remove low-quality cells and potential doublets. Cells were retained if they met the following criteria: (1) total UMI counts between 1,000 and 100,000, (2) number of detected genes ≥ 500, and (3) mitochondrial gene content < 20%. Genes expressed in fewer than 3 cells were excluded from downstream analysis.

### Data Integration and Normalization

To account for technical variation between samples, we employed SCTransform normalization with integration using Seurat. Each sample was individually normalized using SCTransform with mitochondrial gene percentage regressed out to remove technical confounders. Integration features (2,000 genes) were selected, and integration anchors were identified between samples. The integrated dataset was generated using the SCT normalization method, and an additional quality control step removed any remaining cells with mitochondrial content ≥ 20%. Variable features for dimensionality reduction were identified from the RNA assay using variance stabilizing transformation, selecting the top 2,500 highly variable genes.

### Dimensionality Reduction and Clustering

Principal component analysis was performed on the integrated SCT-normalized data using the 2,500 highly variable genes. The first 10 principal components were selected for downstream analysis based on inspection of the elbow plot.

Graph-based clustering was performed using the Louvain algorithm implemented in Seurat. A k-nearest neighbor graph was constructed using the first 10 principal components, and clustering was performed at a resolution of 0.15 to identify major cell populations while avoiding over-fragmentation. This resolution yielded 5 distinct clusters corresponding to biologically interpretable cell states.

For visualization, Uniform Manifold Approximation and Projection (UMAP) was performed using the first 10 principal components with 25 nearest neighbors.

### Cluster Annotation

Clusters were annotated based on expression of canonical cell-type markers. Cluster identities were assigned as follows: Cluster 0 (Regenerative) expressed high levels of *Clu* and *Msln*; Cluster 1 (Stem) expressed *Lgr5* and *Olfm4*; Cluster 2 (Secretory) expressed *Muc2* (goblet cells), *Lyz1* (Paneth cells), and *Chga* (enteroendocrine cells); Cluster 3 (Enterocyte) expressed *Alpi* and other enterocyte markers; and Cluster 4 (Stress response) was characterized by stress-related gene expression signatures.

### seqFISH panel design and quality control

The panel was designed based on a scRNA sequencing dataset of the intestinal monolayer that we had previously collected. Unsupervised clustering of the dataset yielded five main clusters, corresponding to Enterocyte, Stem, Progenitor, Regenerative Response, and Secretory. We chose the top 20 highly variable genes from each cluster and supplemented the resulting list with relevant genes curated from the literature. Probes for each gene were designed by Spatial Genomics for compatibility with the GenePS machine. The final panel targeted 140 genes of interest.

### seqFISH sample preparation and analysis

We prepared and ran the samples using the manufacturer’s protocol. We air-dried the sample and incubated it in a proprietary clearing solution for 10 minutes. We then removed the clearing solution, rinsed in ethanol, and air-dried. We denatured primary probes at 90 °C for 3 minutes. At this point, we applied the primary probe solution and placed a parafilm sheet on top of the coverslip to ensure the probe solution completely covered the coverslip. We then incubated the sample at 37 °C overnight in a hydration chamber. The next day, we washed the sample with a primary wash buffer and then assembled the flow chamber. Once the flow chamber was assembled, we stained the nuclei using rinse buffer with DAPI and then washed the sample several times with rinse buffer. We then loaded the sample into the GenePS machine for rounds of secondary hybridization and readout.

### Data processing, segmentation, and spot assignment

Spots for each region of interest were manually thresholded and subsequently decoded using Spatial Genomics’ SG Analysis Program Software v0.6.8. Nuclei were segmented using Spatial Genomics’ analysis software. We then assigned transcripts to nuclei using the custom-built SGObject package with an expansion of 30 pixels around each nucleus. For expanded nuclei that overlap, transcripts were assigned to the nearest nucleus.

### HCR-FISH (Hybridization Chain Reaction Fluorescence In Situ Hybridization)

#### Sample Fixation

Intestinal organoid monolayers cultured in 96-well plates were fixed for HCR-FISH analysis. The culture medium was aspirated, and the cells were washed twice with PBS. Samples were fixed with 4% formaldehyde in PBS for 10 minutes at room temperature, followed by two washes in PBS. Fixed samples were stored in 70% ethanol at 4°C for a minimum of 12 hours before hybridization.

#### Probe Hybridization

Fixed samples were rehydrated through two washes with 2× SSC buffer, then pre-hybridized in 30% hybridization buffer (30% formamide, 5× SSC, 10% dextran sulfate, 0.1% Tween-20, 9 mM citric acid, pH 6.0, 50 μg/mL heparin, 1× Denhardt’s solution) for 30 minutes at 37°C. Probe pools were diluted to 1.2 pmol per probe set in 30% hybridization buffer. Samples were incubated with probe solution for 6-16 hours at 37°C.

#### Signal Amplification

Excess probes were removed through four washes with pre-warmed 30% wash buffer (30% formamide, 5× SSC, 9 mM citric acid, pH 6.0, 0.1% Tween-20, 50 μg/mL heparin) at 37°C for 5 minutes each, followed by two washes with 5× SSCT (5× SSC, 0.1% Tween-20) at room temperature. Samples were pre-amplified in amplification buffer (5× SSC, 0.1% Tween-20, 10% dextran sulfate) for 30 minutes at room temperature.

Fluorescently labeled hairpin amplifiers (H1 and H2) were snap-cooled by heating to 95°C for 90 seconds, followed by 30 minutes of cooling to room temperature in the dark. Hairpins were diluted in amplification buffer and added to samples for overnight incubation (12-16 hours) at room temperature in the dark.

#### Post-amplification Processing

Excess hairpins were removed through sequential washes: three times with 5× SSCT containing DAPI (1:1000 dilution), followed by two washes with 5× SSCT alone, each for 5 minutes at room temperature. Processed samples were stored at 4°C protected from light until imaging.

### Computational Methods

#### Expected Zone

Following the measurement of gene expression densities in the monolayer based on erosion steps from the edge, we obtain gene expression profiles of all the panel genes as a function of distance from the edge. For each cell, for each erosion iteration, which we define as a zone, we calculate the cosine similarity in expression between the cell’s gene expression profile and the gene expression profile of the zone. Once normalized, this can be considered a discrete distribution of the probability that a cell matches each zone. Therefore, we can calculate the expectation of the zone for each cell.

Mathematically, each cell *j*, *c_j_* ∈ ℝ*^d^* where *d* = 6 is the number of genes measured and used for zone mapping. The genes used for zone mapping are: *Ada*, *Apoa4*, *Apoa1*, *Aldob*, *Alpi*, and *Sis*. The transcript density profiles, *T* ∈ ℝ*^dXm^* where *m* is the number of zones resulting from measuring the transcript densities and binning. In our case m = 11, rings 4-15 (inclusive). The transcript density profile for each zone *t_i_* ∈ ℝ*^d^*, *i* ∈ {1,2, . . *m*}. We computed the probability over zones as: *P*(*t_i_*|*c_j_*) = *cosine similarity* (*t_k_*, *c_j_*)/*∑_k_*_∈{1,.,*m*}_*cosine similarity* (*t_k_*, *c_j_*) Therefore, the expected zone of the cell *j* is: *E*(*t*|*c_j_*) = *∑_i_*_∈{1,2,..*m*}_*P*(*t_i_*|*c_j_*) ∗ *i*

#### Zone Confusion

Given the probability over zones for each cell, we measure zone confusion as the entropy of the distribution.

#### Gene Zone Contribution

For each zone confusion range, a random subset of 10 cells’ normalized expression of each zone mapping gene is shown (expression divided by its sum). The zone mapping genes are shown in order of expression along the villus; therefore, cells with higher zone confusion are expected to express genes related to different zones, whereas cells with lower zone confusion tend to express fewer genes related to different zones simultaneously.

#### Cell–Cell Communication CellphoneDB Analysis

To infer ligand–receptor mediated cell–cell interactions, we applied CellPhoneDB (v0.0.5) on the annotated single-cell transcriptomic dataset derived from intestinal 2D monolayer culture. The analysis was performed using the statistical method of CellPhoneDB ^23^, which computes ligand–receptor pair enrichment across all pairs of cell clusters.

Cells were first grouped by their assigned cell type cluster identities from prior unsupervised clustering. We filtered the expression matrix to include only genes expressed in at least 10% of the cells in a given cluster. For each pair of clusters, CellPhoneDB tested all curated ligand–receptor pairs from its internal database to determine statistically significant interactions based on random permutation of cluster labels (N = 1000). Interactions with a p-value < 0.05 were considered significant.

#### Monolayer computational erosion analysis

To analyze gene expression patterns as a function of distance from the monolayer edge, we created a binary mask delineating the cellular area by performing morphological operations on individual nuclei masks obtained from Cellpose segmentation. We then iteratively eroded this mask inward from the tissue boundary using 4.6 μm intervals. Erosion rings were defined by calculating the difference between consecutive eroded masks, resulting in 20 concentric rings. RNA spot density was calculated for each gene within each ring. Rings 4-15 were used for subsequent analysis as they contained sufficient transcript counts for reliable quantification, while rings 1-3 (nearest the edge) and 16-20 (innermost regions) had sparse had sparse transcript counts

### L-Metric Calculation

#### Overview

We developed a novel metric, termed the L-metric, to quantify the directional association between pairs of genes in single-cell expression data. The L-metric captures asymmetric relationships between gene expression patterns by measuring the deviation of observed co-expression from a null model of independent expression.

#### Data Preprocessing

For each gene pair (gene₁, gene₂), we first filtered cells to retain only those with expression counts ≥2 in at least one of the two genes. This threshold ensures robust statistical estimates while removing uninformative zero-zero expression pairs that dominate sparse single-cell data.

#### Rank-Based Ordering

Following filtering, we created a two-column dataframe containing the expression values for both genes. Cells were then sorted in descending order based on gene₁ expression levels. This rank-based ordering forms the foundation for evaluating how gene₂ expression varies as a function of gene₁ expression rank.

#### Cumulative Distribution Analysis

We computed cumulative sums for both genes across the rank-ordered cells:

- **Observed distribution**: Cumulative sum of gene₂ values in their actual order after sorting by gene₁
- **Null distribution**: Cumulative sum assuming equal distribution of gene₂ expression across all cells (total expression/number of cells)
- **Positive correlation bound**: Cumulative sum with gene₂ values sorted in descending order
- **Negative correlation bound**: Cumulative sum with gene₂ values sorted in ascending order

#### Area-Under-Curve Calculation

The L-metric is derived from the normalized difference between areas under cumulative distribution curves. Using the trapezoidal rule, we calculated:

- A_observed_: Area under the observed cumulative distribution
- A_null_: Area under the null (equal) distribution
- A_positive_: Area under the positive correlation bound
- A_negative_: Area under the negative correlation bound

All areas were normalized by the product of the final cumulative sums of both genes.

#### L-Metric Definition

The raw L-metric is defined as:

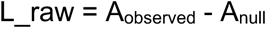

The normalized L-metric is calculated as:

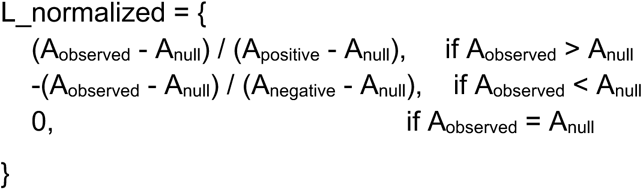

This normalization constrains the L-metric to the range [-1, 1], where:

- Values near +1 indicate a strong positive association (gene₂ expression increases with gene₁ rank)
- Values near -1 indicate a strong negative association (gene₂ expression decreases with gene₁ rank)
- Values near 0 indicate no directional association

#### Pairwise L-Metric Matrix Construction

We computed L-metrics for all ordered gene pairs using permutation analysis. For n genes, this yielded an n × n asymmetric matrix where L(gene₁, gene₂) ≠ L(gene₂, gene₁) in general, capturing the directional nature of gene relationships.

## Data availability

Processed data is available with the paper and on Github: https://github.com/nitzanlab/Resilience-of-spatial-structure-in-intestinal-organoids-revealed-by-spatial-transcriptomics

Raw data is available upon request to the authors.

## Code availability

Code for analysis and figure generation are available on GitHub: https://github.com/nitzanlab/Resilience-of-spatial-structure-in-intestinal-organoids-revealed-by-spatial-transcriptomics

## Acknowledgments

We thank Shalev Itzkovitz for motivating insights and scientific input. We thank Ning Li and Chris Lengner for experimental support and for supplying the H2B-GFP organoid line. We thank Maayan Levi for valuable advice and for assistance extracting the WT organoids. We thank Allon Klein and Ofer Feinerman for fruitful scientific exchanges. We thank members of the Raj and Nitzan labs for helpful feedback on the work.

This project has been made possible in part by grant 2023-332391 from the Chan Zuckerberg Initiative DAF, an advised fund of Silicon Valley Community Foundation. A.R. acknowledges support from the Samuel Waxman Cancer Research Foundation, and the Templeton Foundation (63532). M.N. acknowledges support by the EMBO Young Investigator Programme, the Abisch-Frenkel Foundation, and the European Union (ERC, DecodeSC, 101040660). Views and opinions expressed are, however, those of the author(s) only and do not necessarily reflect those of the European Union or the European Research Council. Neither the European Union nor the granting authority can be held responsible for them.

## Author contributions

Y.H. and A.R. conceived and designed the project. M.E. and M.N. designed new analytical tools. Y.H. designed and performed experiments. Y.H. and M.E. performed data analysis under the supervision of A.R. and M.N. P.B. produced the intestinal monolayer single-cell RNA-seq dataset and provided valuable support during initial phases of the project. Y.H. and M.E. wrote the manuscript with input from all authors. All authors read and approved the final manuscript.

## Declaration of Generative AI and AI-Assisted Technologies

During the preparation of this work, the authors used ChatGPT and Claude to generate and improve code for analyses. The authors reviewed and edited the content and take full responsibility for the content of the manuscript.

## Supplementary Figures

**Supplementary Figure 1:**
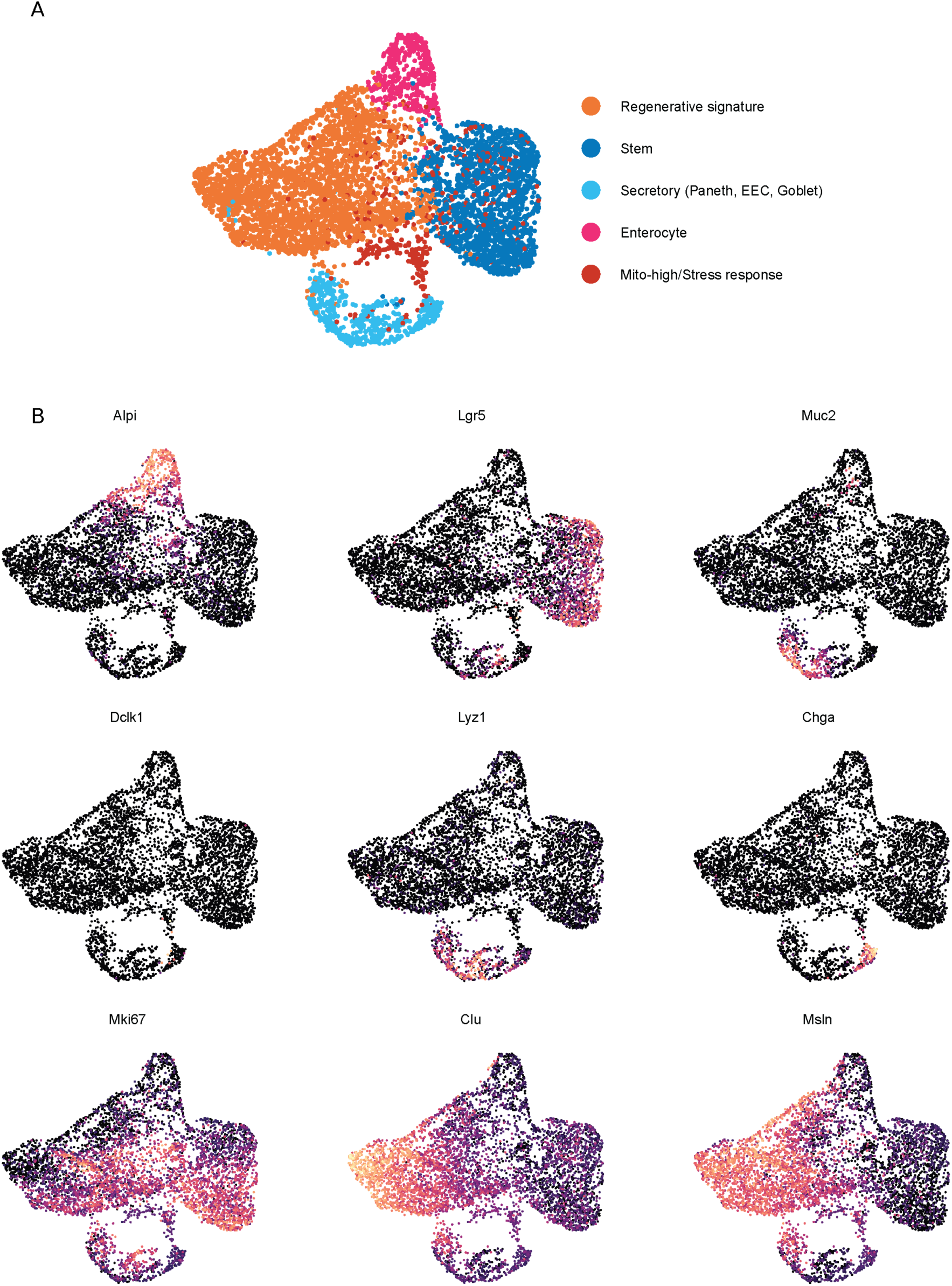
Intestinal monolayer cell type clustering based on single-cell RNA sequencing data. a). UMAP embedding of single-cell RNA-seq profiles from 2D murine intestinal epithelial monolayers. Cells are colored by Seurat clusters (resolution 0.15) and labeled by cluster ID. b). UMAP FeaturePlots showing normalized SCT expression of marker genes across single cells from 2D murine intestinal epithelial monolayers.

**Supplementary Figure 2:**
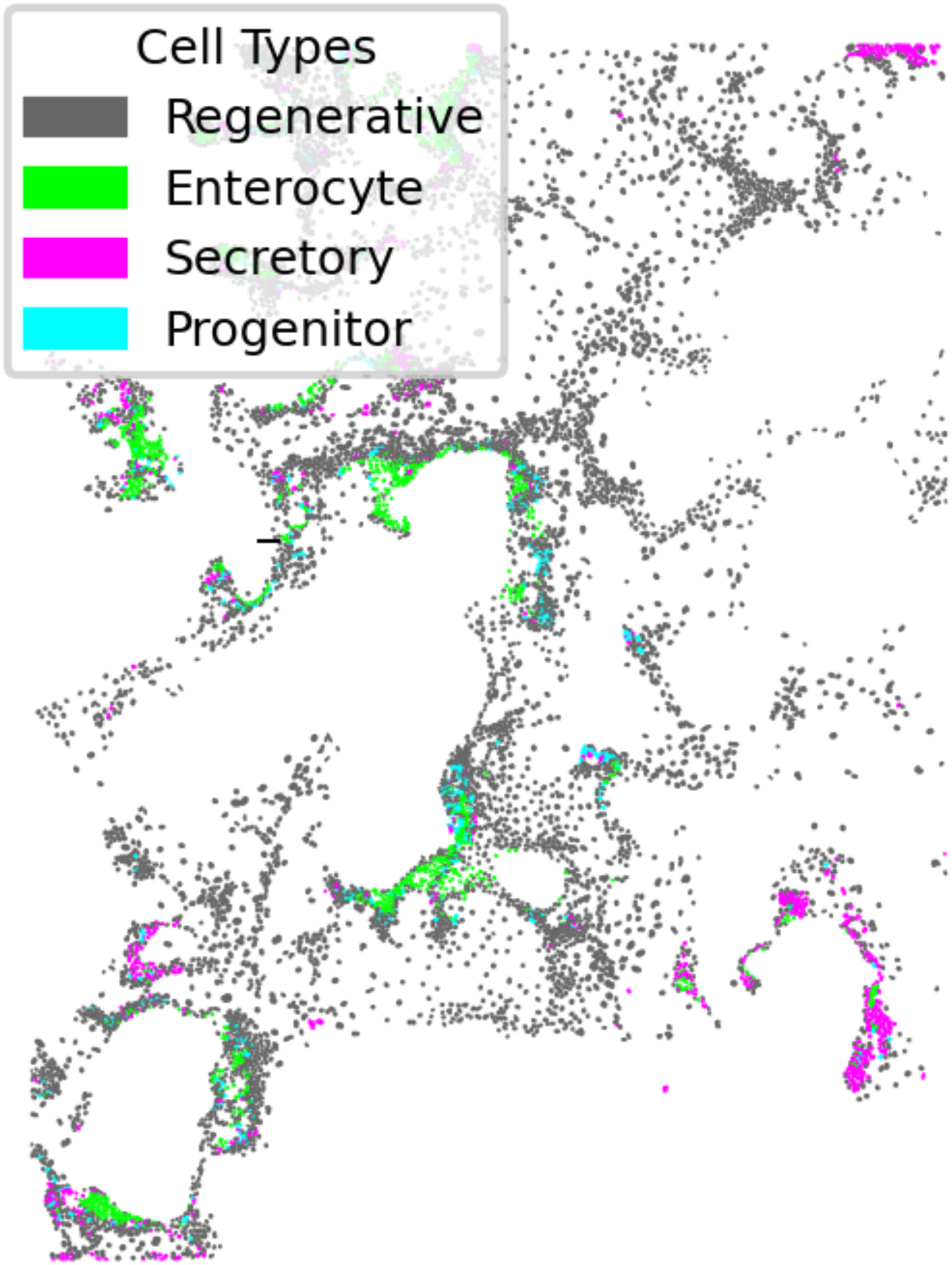
Intestinal monolayers colored by cluster identity.

**Supplementary Figure 3:**
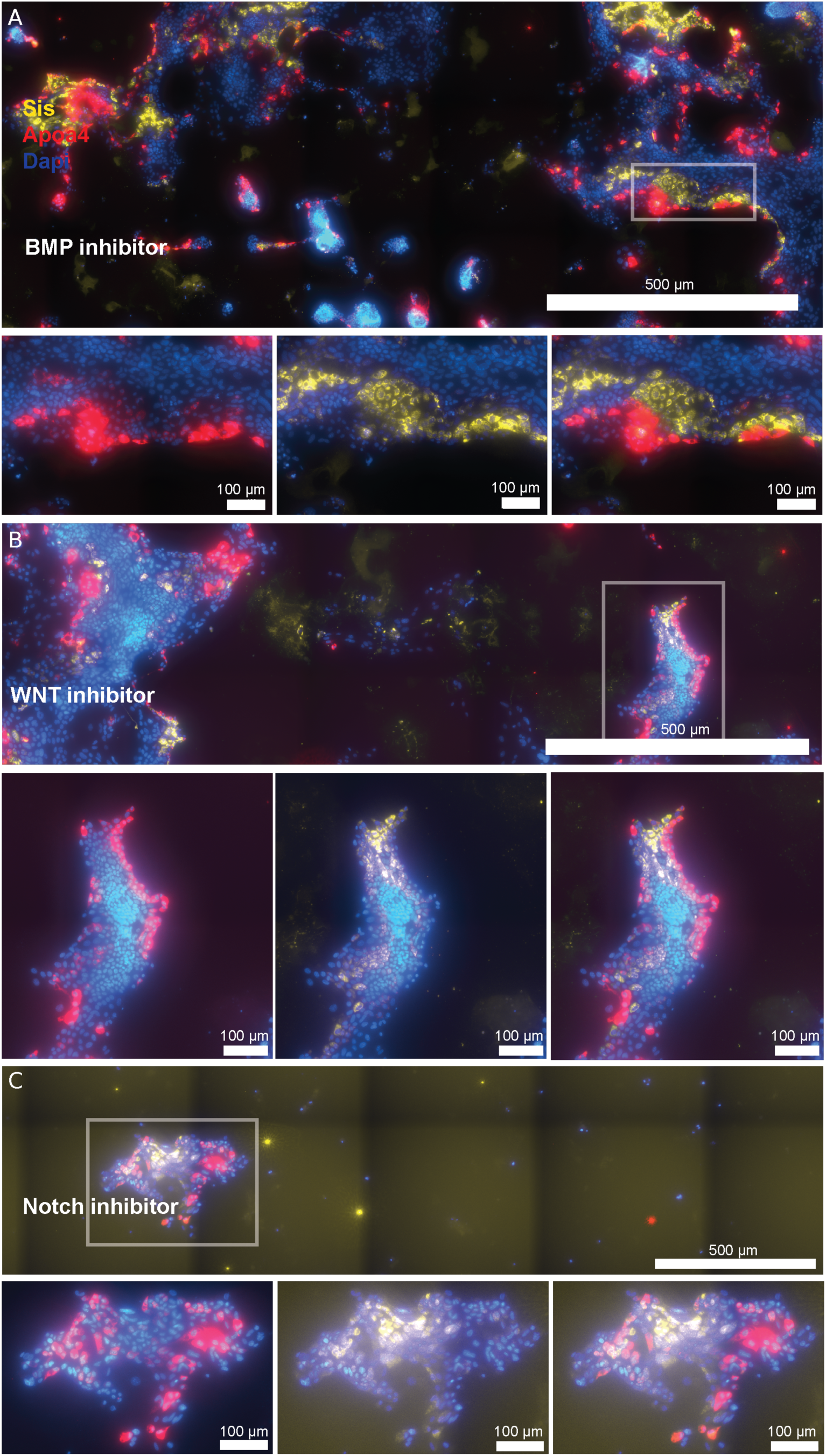
Extended perturbation experiment results. Images of intestinal monolayers cultured with various pharmacological agents and profiled for spatial mRNA expression of *Sis* (highly expressed at the villus bottom) and *Apoa4* (highly expressed at the villus tip). a) BMP inhibitor, LDN b) WNT inhibitor, IWP-2 c) Notch inhibitor, R0

**Supplementary Table 1:**
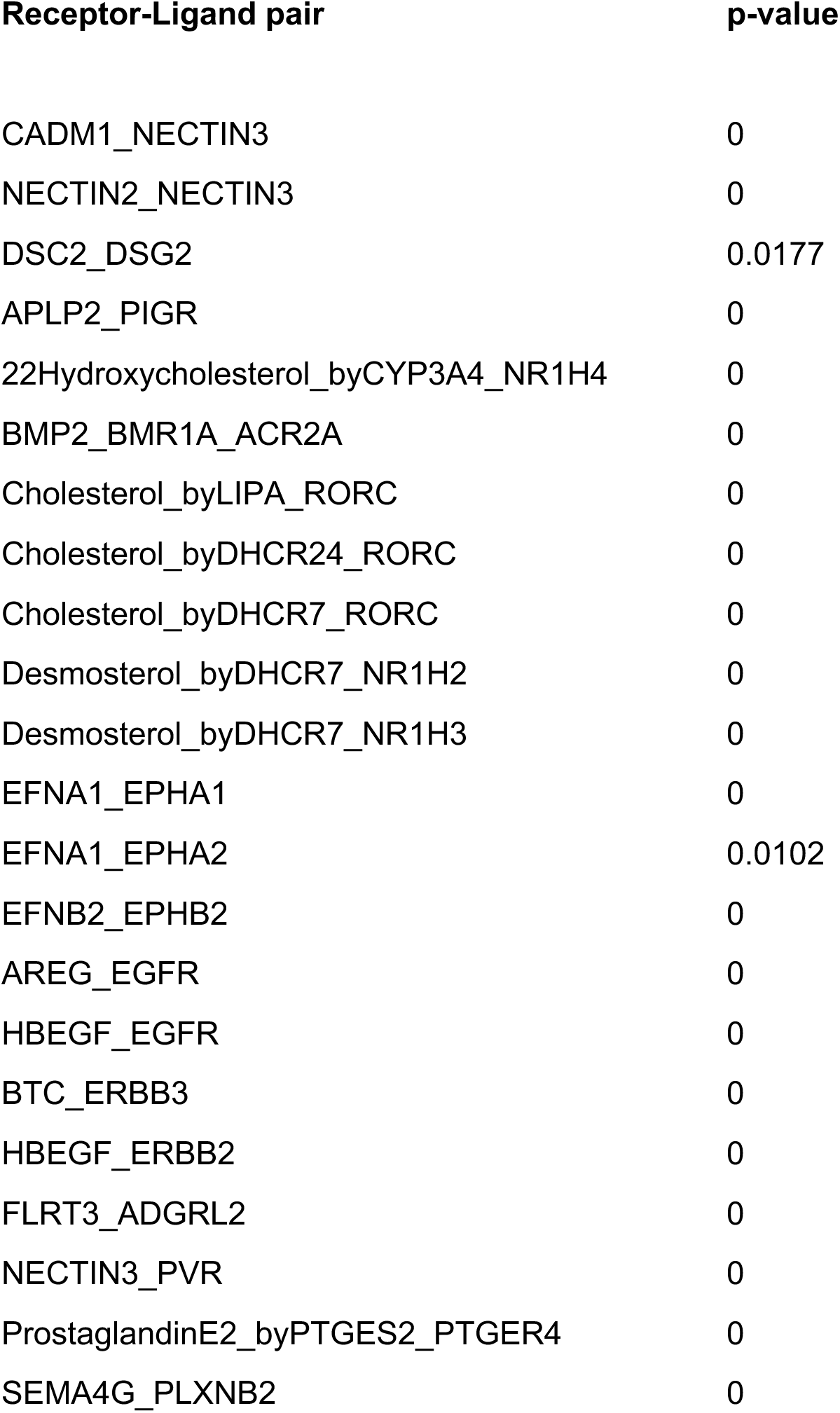
Receptor-Ligand pair candidates in the enterocyte population identified by cellphoneDB.

**Supplementary Figure 4:**
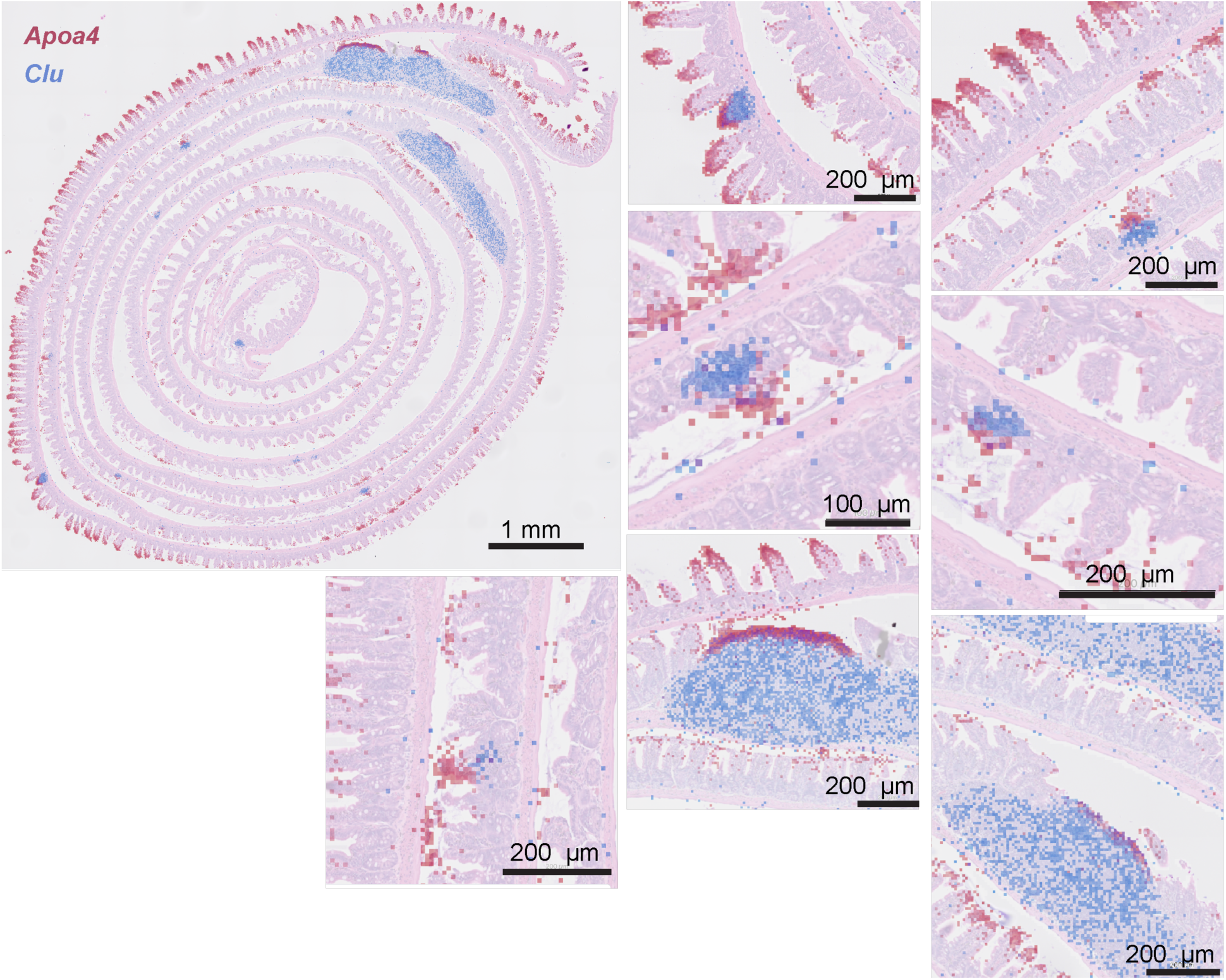
Truncated villi express villus top markers. Mouse intestine cross-section with spatial expression patterns of *Clu* (blue) and *Apoa4* (red) captured by Visium HD (10X Genomics). Truncated injured villi are identified by high *Clu* expression.

